# Oxytocin involves in chronic stress-evoked melanoma metastasis via β-arrestin 2-mediated pathways

**DOI:** 10.1101/494484

**Authors:** Haoyi Ji, Na Liu, Jing Li, Dawei Chen, Qian Sun, Yingchun Yin, Yanli Liu, Bing Bu, Xiaoyang Chen, Jingxin Li

## Abstract

Stress is associated with an increased risk of lung metastasis in melanoma. However, the underlying mechanism is elusive. Oxytocin (OXT), a neurohormone produced by the hypothalamus, plays a vital role in laboring induction and lactation. Emerging evidence suggests that OXT also regulates human emotions, social cognition, social behaviors and stress-related disorders. **Here, we reported that** a significant up-regulation of oxytocin receptors (OXTRs) was observed in malignant melanoma. The activation of oxytocin receptors (OXTRs) dramatically promoted migration, invasion and angiogenesis but not the proliferation of melanoma cells *in vitro* and *in vivo* via β-arrestin 2-dependent ERK-VEGF/ MMP-2 pathway. Next, chronic restraint stress significantly elevated the plasma level of OXT. Notably, 21 days chronic restraint stress facilitated lung metastasis of melanoma and reduced overall survival in mice, which were largely abrogated by knocking down either OXTR or β-arrestin 2. These findings provide evidence that chronic stress hormone-OXT promotes lung metastasis of melanoma via a β-arrestin 2-dependent mechanism and suggest that OXT, a novel pro-metastasis factor, is a potential therapeutic target for melanoma.

**Importance of the Study:** Oxytocin, a neuropeptide produced in the hypothalamus that is involved in lactation and parturition, play a vital role in chronic stress-evoked melanoma metastasis via β-arrestin 2-mediated ERK signaling pathway, which provides a potential therapeutic target for melanoma.

## Introduction

Melanoma is the most deadly type of skin cancer and its incidence rate is rising worldwide. Presently, there are no effective therapies to prevent melanoma metastasis. As the only Food and Drug Administration (FDA)-approved melanoma adjuvant therapy, interferon-alpha only showed a meager 5-year overall survival benefit of 1%– 3% [1]. Despite recent advances in immunotherapy, most metastatic melanoma patients still die from their own disease, with an average survival time of 23 months [2, 3]. For patients with advanced melanoma, due to its poor clinical prognosis and lack of effective treatment, it is necessary to develop and implement new methods to prevent metastatic recurrence of this cancer. A growing body of literature suggests that stress can affect cancer progression. Chronic stress increases the spread of metastatic cancer cells from primary breast tumors to the lungs [4].

Stress hormones promote EGFR inhibitor resistance in the treatment of non-small cell lung cancer[5]. If cancer patients are socially isolated or separated from their spouses, their survival rate decreases markedly [6, 7]. Instead, social support can improve the prognosis of cancer patients [8]. Chronic restraint stress promotes melanoma lung metastasis by inhibiting the immune system [9, 10]. However, there are no rational explanations for the poor outcomes of stress-associated melanoma patients so far. Oxytocin (OXT), a neurohormone produced by the hypothalamus, plays a vital role in laboring induction and lactation. Emerging evidence suggests that OXT also regulates human emotions, social cognition, social behaviors [11] and stress-related disorders [12]. Hormone OXT also belongs to stress hormones and is released in response to several stress stimuli [13]. After exposed to repeated restraint stress, the concentrations of OXT in plasma was significantly increased [14]. Therefore, we hypothesized that stress hormone-OXT involves the stress-promoted melanoma pulmonary metastasis. Recently, an additional new role of OXT has been identified in neoplastic pathology. In tumor, OXT acts as a grow regulator, through the activation of a specific G-coupled transmembrane receptor, the oxytocin receptor (OXTR) [15]. Although OXT causes tumorigenesis, the role of OXT in malignant melanoma is unclear, and whether OXT signals will be a target for melanoma treatment remains unclear. The purpose of this study was to investigate the role of OXT in the lung metastasis of stress melanoma. Herein, we demonstrated that OXTR activation increased melanoma cells migration and invasion *in vitro*, and promoted lung metastasis of melanoma *in vivo by* stimulating ERK1/2 phosphorylation and MMP-2 and VEGF production, which was abrogated with knockdown of either OXTR or β-arrestin 2. The findings of present study shed some light on the mechanism of the poorer outcomes of the stress-related promotion of melanoma metastasis, suggesting OXT targeting might be a new strategy for melanoma therapy.

## Materials and methods

### Collection of malignant melanoma samples

All specimens were confirmed as malignant melanoma by pathological diagnosis. All patients involved in this study received no radiation therapy or chemotherapy before surgical resection. The study included 7 females and 8 males, ranging in age from 41 to 71 years with a mean age of 61 years. The clinicopathological information is summarized in Supplemental table 1. Tissues were snap frozen in liquid nitrogen, and then stored at −80°C before use.

### Measurement of OXT

One week after stress, mice (n=8) were anesthetized, blood was collected using a pre-chilled 1ml EDTA tubes from the caudal vein. Plasma (200μl) was isolated by centrifugation at 1800 rcf, 15 minutes at 4°C. OXT contents were quantified in duplicates using a commercial OXT ELISA kit (Enzo Life Sciences, NY, USA).

### Cell culture and reagents

The human melanoma cancer cell line A375 and the murine melanoma cell B16 were purchased from China Center for Type Culture Collection. A375 cells were cultured in DMEM (Hyclone Co., Logan, UT, USA) supplemented with 10% (v/v) FBS (Gibco), 100 U/mL penicillin and 100 μg/ml streptomycin. The B16 cells were cultured in RPMI 1640 (Hyclone Co) supplemented with 10 % FBS, 100 U/ml penicillin and 100 µg/ml streptomycin. All cell lines were propagated at 37°C in an atmosphere containing 5% CO_2_. Primary antibodies for OXTR, MMP2 and VEGF were purchased from Abcam (Austin, TX), those for β-arrestin 1, β-arrestin 2, Erk and p-ErK were purchased from CST (Beverly, MA), that for b-actin was from Sigma-Aldrich (St Louis, MO). OXT was purchased from Sigma-Aldrich (St Louis, MO).

### Xenograft murine model

For orthotopic mouse tumor model, 5×10^5^ B16 melanoma cells were injected *s.c*. into right hind flank of 6-week-old male C57BL/6 mice to develop melanoma. After one week of inoculation, mice with approximate tumor size were divided into two groups and administered intraperitoneally with normal saline or OXT (0.1 mg/kg/d) for 2 weeks. The tumor volume was calculated according to the formula (length × width^2^)/2 [16]. Three weeks after tumor inoculation, they were then sacrificed for evaluating primary tumor size and weight. For the chronic stress model, we used a previously described and well-defined restraint stress procedure to induce symptoms of stress[17]. In this model, mice are securely restrained in a movement-restricted space for 21 days to 2 hours of restraint in an acrylic cylindrical animal restrainer with holes that permit the restrainer to be adjusted according to the size of the subject. The restrainer allows unlimited breathing but restricts the movement of the limbs. After being restrained, the mice were returned to their home cage and given food and water. In all experiments, 5×10^5^ B16 cells were injected via tail vein (to study metastasis) in mice 1 week after the restraint stress procedure began. After 14 days, the animals were killed by cervical dislocation, and the numbers of tumor nodules of lung metastases were noted by researchers blinded to the groups. The lungs were harvested for immunohistochemical analysis. Following the same procedure, the survival of the mice was observed.

### Plasmid construction and generation of stable clones

The siRNA sequence targeting mouse β-arrestin 2 5′-AAGGACCGGAAAGUGUUCGUG-3 is reported elsewhere[18]. β-arrestin 2 shRNA (shβ-arr2) plasmid was constructed by Genechem (Shanghai, China). OXTR shRNA plasmid (shOXTR) and control plasmid were purchased from Genechem (Shanghai, China). Transfections were performed in 24-well culture plates using Lipofectamine 2000 (Invitrogen, Carlsbad, CA, USA) according to the manufacturer’s instructions. Two days after transfection, B16 cells were selected by 1μg/mL puromycin (Sigma-Aldrich, StLouis, MO). The expression of β-arrestin 2 and OXTR after transfection was detected by Western blot.

### Western blot and immunocytochemical analysis

For Western blot analysis, cells were lysed in lysis buffer [50 mmol/L Tris (pH 7.5), 100 mmol/L NaCl, 1 mmol/L EDTA, 0.5% NP40, 0.5% Triton X-100, 2.5 mmol/L sodium orthovanadate, 10 μl protease inhibitor cocktail, 1 mmol/L phenylmethylsulfonyl fluoride] for 30 min at 4°C. Total proteins were fractionated by SDS-PAGE and transferred onto nitrocellulose membrane. The membranes were then incubated with appropriate primary antibodies (OXTR, MMP2,VEGF, β-arrestin 1, β-arrestin 2, Erk and p-ErK), followed by incubation with a secondary HRP-conjugated antibody. For histopathologic analysis, lung samples were fixed in 4% paraformaldehyde immediately after being harvested and then paraffin-embedded. Lung samples were subjected to antigen retrieval treatment by microwave prior to blocking with sequential peroxidase and goat serum block. After incubation for the diluted primary antibody, a secondary HRP-conjugated antibody was used. For protein levels, the slides were stained with respective antibodies and staining intensity assessed semi-quantitatively. In short, 5 random fields were chosen per slide and scored from 0 to 4 on intensity, and 0% to 100% on distribution of positive staining of tumor tissues. The final results per group are presented in the accompanying graph with representative pictures for each group.

### Gelatin zymography analysis

The effects of OXT on the gelatinolytic activities of MMP-2 were examined by gelatin zymography. A375 cells were treated with various concentrations of OXT for 48 hours in serum-free medium. Serum-free supernatants were then harvested and centrifuged to remove cellular debris. Protein concentrations were determined with the bicinchoninic acid assay protein reagent kit (Sangon, Shanghai, China). An equal amount of protein (20 μg) from each treatment was diluted with the loading buffer and fractionated on 10% SDS / PAGE containing 1 mg/mL gelatin A (Sigma Chemical Co.). After electrophoresis, the gels were stained with 0.1% Coomassie Brilliant Blue and destained with 45% methanol, 10% (v / v) acetic acid until clear bands suggestive of gelatin digestion were present.

### Cell proliferation studies

The proliferation of B16 and A375 cells was assessed by Cell Counting Kit-8 (CCK-8, Beyotime, C0038) assay according to the protocol recommended by the manufacturer. The approximate 2000 cells were seeded in each well of 96-well plates with 100 μL medium. OXT (10^−7^ M) was added to each well. After being incubated for 1, 2, 3 and 4 days, 10 μL CCK-8 solution was added into each well. Cells were then incubated for another 4 h at 37°C. The absorbance was measured by a microplate reader at the wave length of 450 nm. In each experiment, six parallel wells were set up.

### Wound-healing assay

B16 cells were seeded onto a six-well culture plate and cultured to a subconfluent state in complete medium. Cell monolayers were linearly scraped with a P-200 pipette tip (250-μm width). Cells detached from the bottom of the well were mildly aspirated and incubated in serum-free medium containing different concentrations of OXT (0,10-9,10-8,10-7,10-6 mol/L) for 24 hours. The width of the scratch was microscopically monitored at various time points and quantified in terms of the difference between the original width of the wound and the width after cell migration. The percentage of wound closure [(original width-width after cell migration)/original width] was calculated. The width of the wound was measured using Image-Pro Plus 6.0.

### Transwell migration assay

A complementary transwell migration assay was performed by employing a modified Boyden chamber (Corning Costar) containing a gelatin-coated polycarbonate membrane filter (pore size, 8 μm). A total of 2×10^4^ cells in 500 μL of culture medium containing various concentrations of OXT (0, 10^−9^, 10^−8^, 10^−7^, 10^−6^ mmol/L) were added to the upper chamber, and the lower chamber contained culture medium with 10% FBS to stimulate cell migration. The migration assays were incubated for 24 hours at 37°C in 5% CO_2_, and then the cells were stained with crystal violet. Cells on the undersides of the filters were observed under a microscope at a magnification of 200 x and counted.

### Matrigel in vitro HUVEC tube formation assay

The *in vitro* tube formation assay was performed according to the manufacturer’s recommendation. Briefly, growth factor-reduced Matrigel, after being thawed on ice, was plated in pre-cooled 12-well chamber. After solidification of Matrigel (30 min incubation at 37°C), HUVECs were trypsinized and seeded (5×10^4^ cells per well) in each well with 1640 medium alone (control sample) or 1640 supplemented with cell-free culture supernatants from different treatment (1:1, v/v). Each sample was added in duplicate wells. The chambers were then incubated at 37°C for 24 hours. After incubation and fixation phases, the tube formation process involving HUVECs adhesion, migration, differentiation and growth was scored using an inverted phase-constrast microscope and measured using Image-Pro Plus 6.0.

### Statistical analysis

Continuous variables were recorded as mean ± SD and analyzed by Student’s *t* test. For data with non-normal distribution or heterogeneity of variance, median (range) was shown and comparison was examined by Mann-Whitney *U* test or Kruskal-Wallis *H* test. All survival analyses were estimated from Kaplan-Meier curves. The log-rank test was used to assess the significance of univariate survival analyses. Statistical analysis of the results was performed using GraphPad StatMate software (GraphPad Software, Inc, San Diego, CA). The two-sided *P*-values <0.05 was considered statistically significant.

## Results

### The OXTRs were expressed in melanoma cells and significantly upregulated in malignant melanoma compared to controls

Next, we examined whether OXTRs were expressed in melanoma cells. As expected, Western blot analyses of cultured mouse (B16) and human (A375) melanoma cells, human malignant melanoma tissues and the paratumor tissues **(Table 1)** with OXTRs antibody detected the expression of OXTRs (Fig.1A). The melanoma cells contained high level of OXTRs protein. Strikingly, the OXTR protein levels of human melanoma tissue samples were significantly upregulated compared with the paratumor tissues (n=5), which were confirmed by immunochemistry analyses (n=10, Fig.1B). These above results indicate that OXT signaling is in the development of melanoma.

**Fig. 1:**
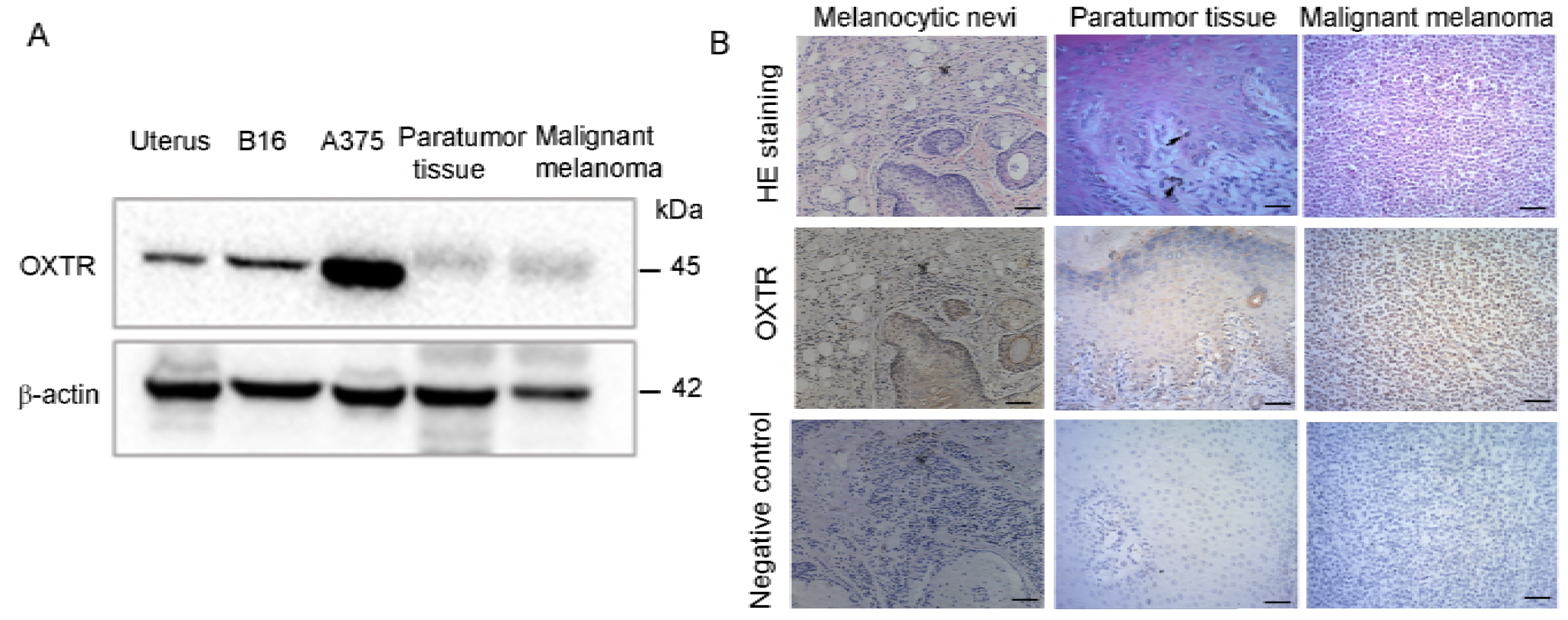
OXTRs were expressed in melanoma cells and significantly upregulated in malignant melanoma. (A) Western blot analyses of cultured mouse (B16) and human (A375) melanoma cells, human malignant melanoma and paratumor tissues with antibodies to OXTR. Uterus: positive control. Notably, the expression of OXTR in malignant melanoma was significantly upregulated compared with paratumor tissues (n=5). (B) Immunohistochemical staining of samples from melanocytic nevi (n=3), malignant melanoma and paratumor tissues showing the expression of OXTR (n=10). The OXTR was overexpressed in malignant melanoma compared to melanocytic nevi or paratumor tissues. Black arrows show Melanocytes.

### OXT promoted migration, invasion and angiogenesis but not proliferation of melanoma cells in vitro and in vivo

Next, we investigated the role of OXT in the migration, invasion and angiogenesis of melanoma cells. The transwell migration assay and wound healing assay revealed that OXT markedly increased the migration and invasion of both B16 mouse melanoma cells and A375 human melanoma cells in dose-response manner (*P* <0.05;Fig.2A-B). It is well known that tumor growth and metastasis depend on angiogenesis [19]. The tube formation assay is one of the most widely used *in vitro* assays to model the reorganization stage of angiogenesis. As expected, the tube formation assay revealed that the supernatants of A375 melanoma cells treated with OXT markedly promoted HUVECs tube formation *in vitro* (*P* < 0.05;Fig. 2C). To verify whether OXT affects the migration and invasion of melanoma cell *in vivo*, we utilized a murine xenograft model of spontaneous metastases of B16 mouse melanoma cells. The administration of OXT largely enhanced the lung metastasis of B16 melanoma cells in C57BL/6 mice (n = 9, *P* < 0.05;Fig.3A) and reduced the survival in mice with melanoma lung metastasis compared with the PBS control group (n = 8, *P* =0.056; Fig.3B). However, atosiban, an antagonist of OXTR had no effect on the lung metastasis and the survival rate (n = 8-9, *P* >0.05; Fig.3B).

**Fig. 2:**
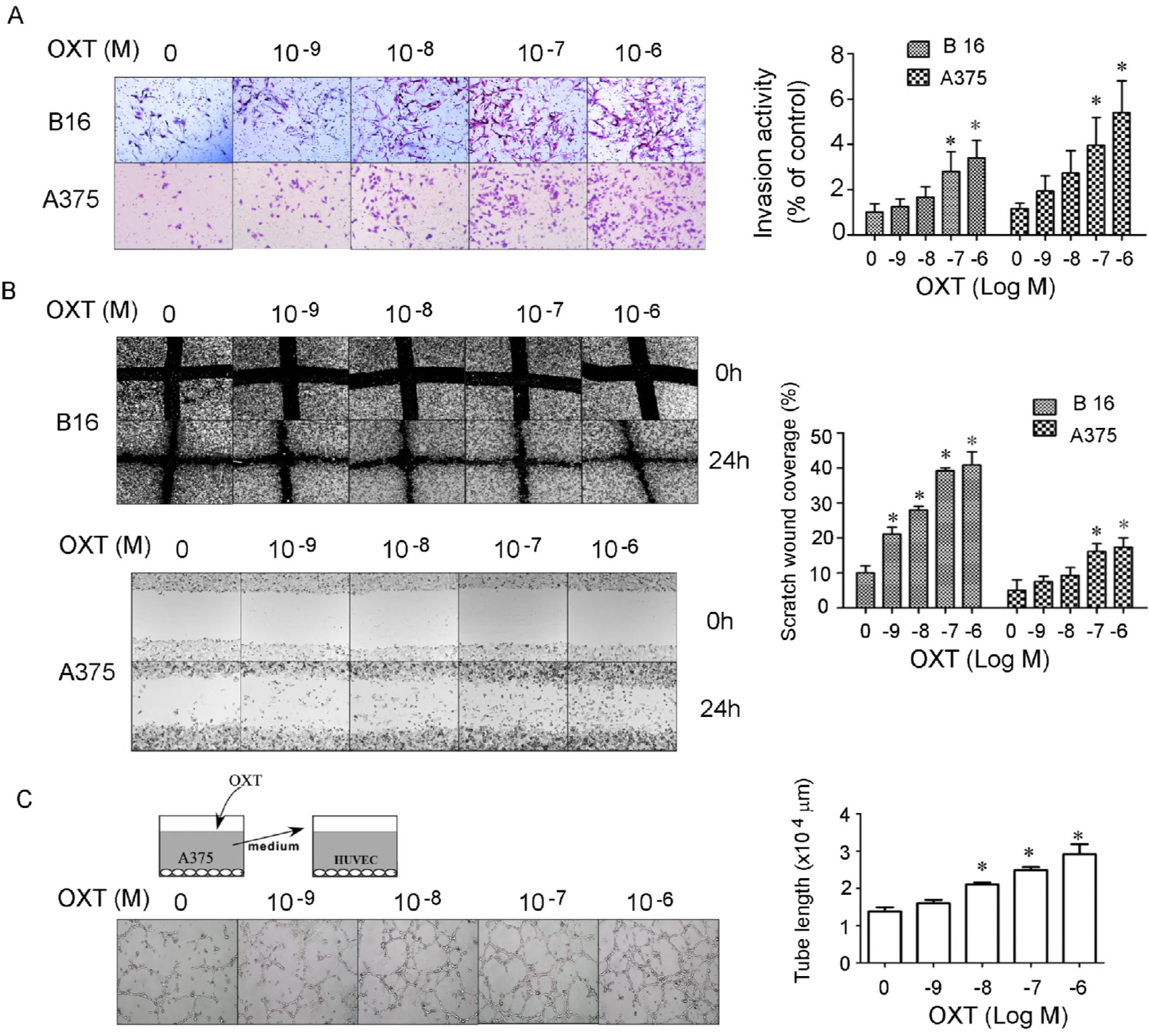
Effect of OXT on melanoma cell migration and angiogenesis activity *in vitro*. (A) B16 and A375 cells were seeded in the upper chamber treated with 0, 10^−9^, 10^−8^, 10^−7^ and 10^−6^ M of OXT and allowed to migrate for 24 hours; then, the migrated cells were stained with crystal violet. The graphs depict the mean ± SD of three independent experiments. *, *P*< 0.05.(B) The effects of OXT on B16 and A375 cells migration ability were plotted as the percentage of wound closure. The graphs depict the mean ± SD of three independent experiments. *, *P* < 0.05. (C) Angiogenesis was measured by *in vitro* tube formation assay. HUVECs were cultured with supernatants of A375 cells treated with serial concentration OXT on Matrigel. The degree of tube formation was quantified by Image-Pro Plus 6.0. The graphs depict the mean ± SD of three independent experiments. *, *P* < 0.05.

To observe whether activation of OXTR affects proliferation in melanoma cells, we treated melanoma cells with OXT or vehicle *in vitro* (*P* > 0.05;Fig.3C), or injected *s.c*. equal numbers of B16 mouse melanoma cells into the right hind flank of host C57BL/6 mice (Fig. 3D) followed by successive injection of OXT (0.1mg/kg/d). B16 is a selective variant cell line obtained from pulmonary metastasis of a melanoma, syngeneic to black C57BL/6 mice [20]. However, OXT did not affect cancer cell proliferation(Fig. 3C) and tumor growth(*n* = 5, *P* > 0.05; Fig. 3E).

**Fig. 3:**
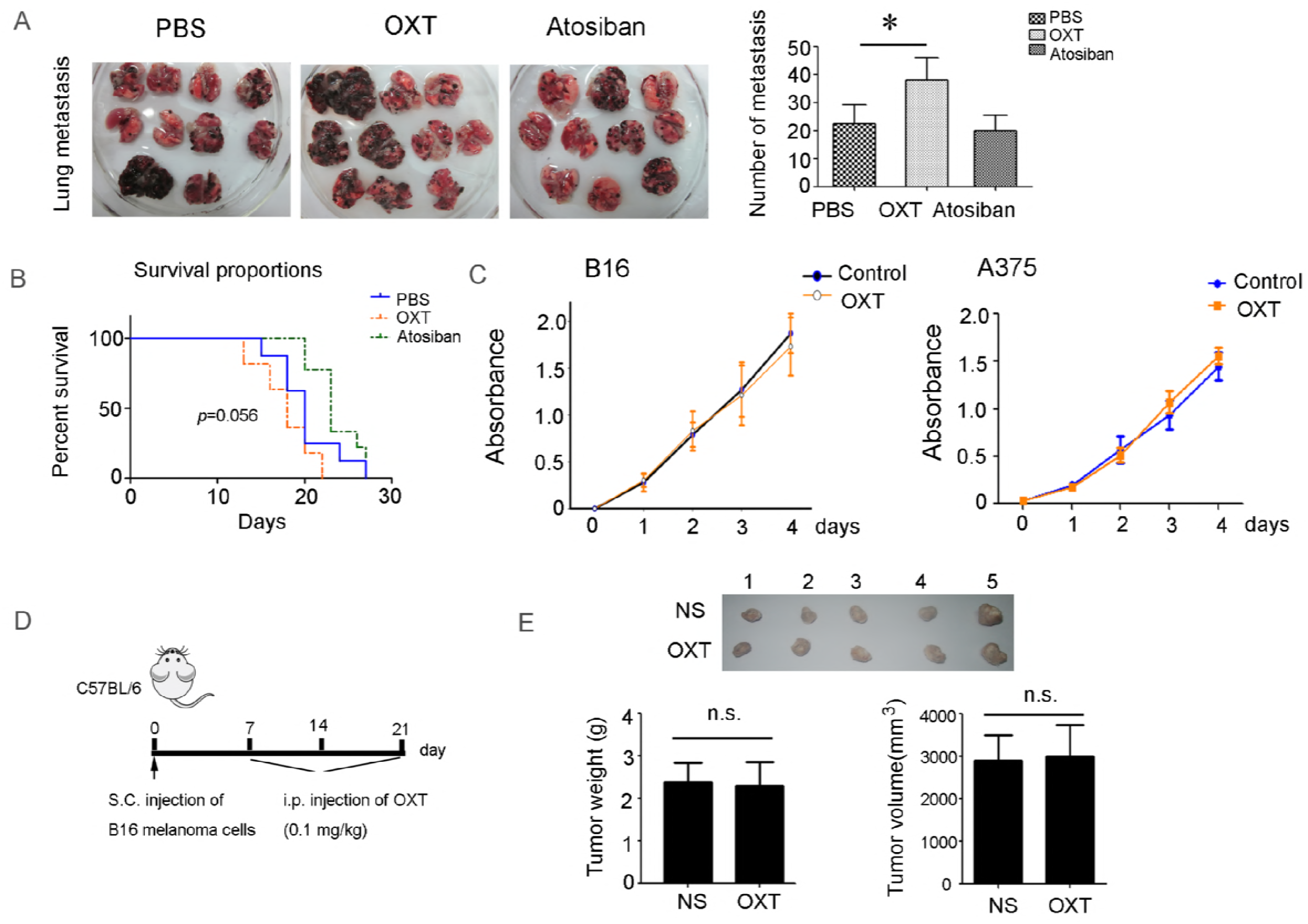
Effect of OXT on melanoma lung metastasis and proliferation *in vivo* and *in vitro*. (A) Three days after B16 cell injection, mice divided randomly into three groups and treatment with OXT, atosiban or PBS for 1 week. 14 days after B16 cell injection, the animals were killed by cervical dislocation and the lungs were harvested. Representative images and the quantification of the metastatic foci of the lungs from mice 14 days after B16 injection (*n* = 9). *, *P* < 0.05. (B) Survival of mice treatment with PBS, atosiban, or OXT followed by *i.v*. injection of B16 cells (*n*= 8). (C) The approximate 2000 cells were seeded in each well of 96-well plates with 100 μL medium. OXT (10^−7^ M) was added to each well. The proliferation of B16 and A375 cells was assessed by Cell Counting Kit-8 assay. After being incubated for 1, 2, 3 and 4 days, 10 μL CCK-8 solution was added into each well. Administration of OXT did not affect the proliferation of melanoma cells *in vitro*. (**D**) Three weeks after injection with 1×10^6^ B16 tumor cells into the inguinal area of host mice, the tumor size or weight were measured. Administration of OXT (0.1mg/kg/d) did not affect the tumor size or tumor weight (E, *n*=5). n.s.: No significance.

### OXT stimulated the production of VEGF and MMP-2 and promoted metastasis via a β-Arrestin2-dependent ERK1/2 activation in melanoma cells

To explore the mechanism of OXT-induced metastasis, firstly we detected the production of VEGF and MMP-2. Western blot assay revealed that OXT dose-dependently promoted the production of VEGF and MMP-2 in both B16 mouse melanoma cells and A375 human melanoma cells, and led to robust ERK1/2 activation (Fig. 4 A). Using gelatin zymography analysis, we further confirmed OXT increased the MMP-2 activities of melanoma cells in dose-dependent manner (Fig.4B). β-arrestins have been shown to bind several signaling proteins, including Src [21], ERK1/2[22-24] and JNK3 [25] in an agonist-dependent manner. To assess the importance of β-arrestin 2 in mediating the G protein independent activation of ERK1/2, we used the siRNA interference to down-regulate the expression of endogenous β-arrestin 2 and examined ERK1/2 activation by OXT. Silencing either OXTR or β-arrestin 2 largely suppressed the expression of OXTR or β-arrestin 2 in melanoma cells (Fig. S1). Down-regulation of β-arrestin 2 expression by siRNA transfection abolished the ERK1/2 activation induced by OXT in B16 mouse melanoma cells (Fig. 4C). Notably, the down-regulation of β arrestin 2 expression by siRNA also inhibited the enhancement of VEGF and MMP-2 activated by OXT treatment (Fig. 4C). This finding indicates that the G protein-independent activation of ERK1/2 evoked by OXT is completely dependent on β-arrestin 2. Because β-arrestin 1 was not expressed in B16 melanoma cells (Fig. S2), these results suggest that β-arrestin 2, not β-arrestin 1, is the major form of β-arrestin mediating OXT-induced G protein-independent ERK1/2 activation.

**Fig. 4:**
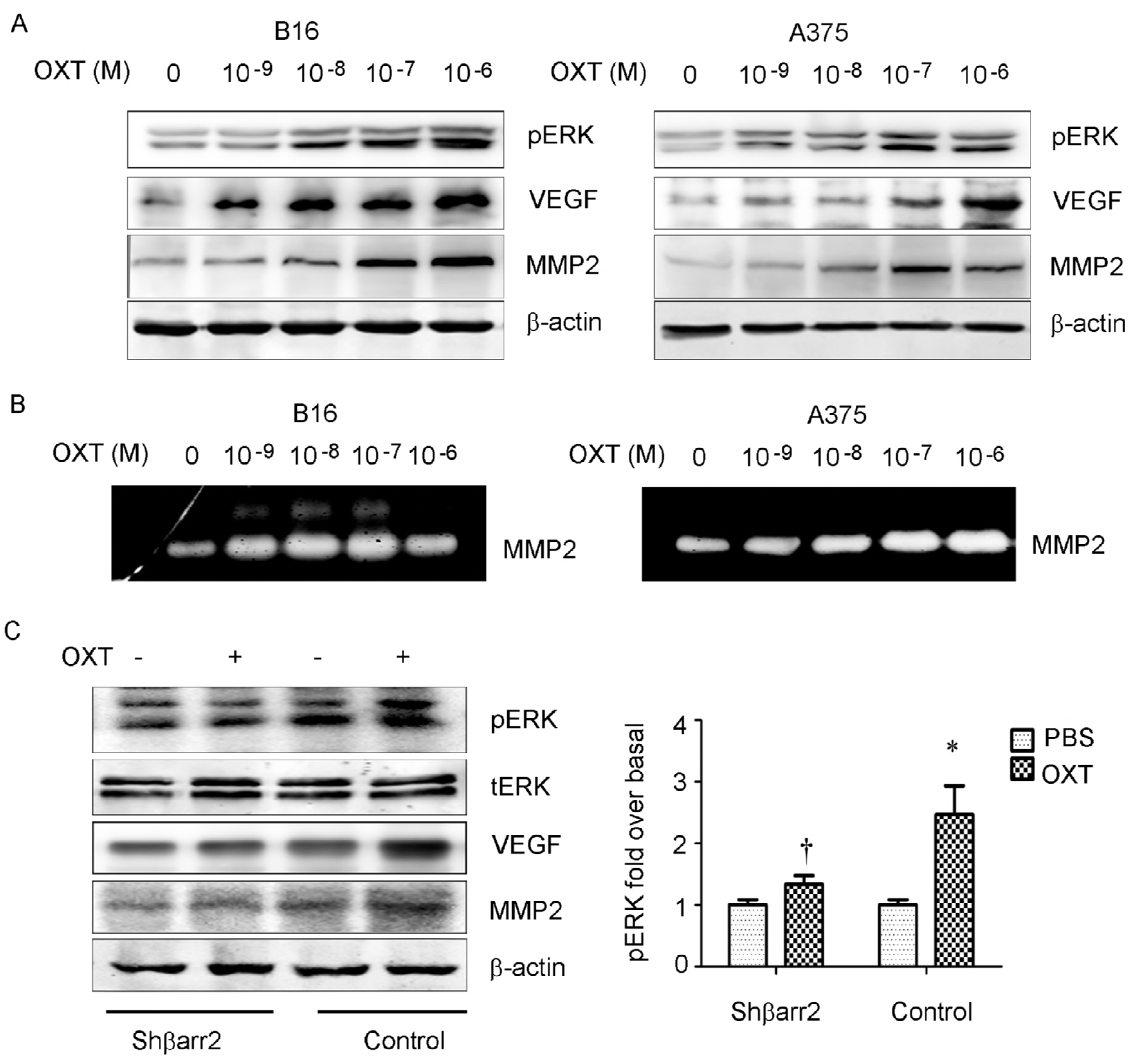
OXT enhances the production of VEGF and MMP2 via a β-arrestin 2-dependent ERK. (A) B16 and A375 cells were treated for 0, 10^−9^, 10^−8^, 10^−7^ and 10^−6^ M of OXT for 48 hours. The cells were lysed and processed on a Western blot and probed for p-ERK and VEGF, MMP2. (B) B16 and A375 cells were treated for 0, 10^−9^, 10^−8^, 10^−7^ and 10^−6^ M of OXT for 48 hours. MMP2 activities in OXT-treated cells were detected using gelatin zymography analysis. (C) Control and β-arr2 down-regulating B16 cells were treated for 0 or 10^−7^ M of OXT for 48 hours. Phosphorylated ERK (p-ERK), total ERK (tERK), MMP2 and MMP9 were measured by Western blot. Mean densitometry from three independent experiments demonstrated that OXT increased p-ERK signaling (* *P*< 0.05), but the response was diminished in β-arrestin 2 down-regulating cells compared with control cells († *P*< 0.05, β-arrestin 2 down-regulating cells receiving OXT compared with control cells receiving OXT).

### Chronic restrain stress promoted melanoma lung metastasis, which were largely abolished by knocking down either OXTR or β-arrestin2

To explore the role of stress in lung metastasis of melanoma cells, we utilized a repeated restrain stress mouse model (Fig.5A). As expected, restrain stress significantly promoted the melanoma lung metastasis (n=8, *P* < 0.05; Fig.5B) and reduced the survival of mice with metastatic cancer (n=8, *P* < 0.05;Fig.5C) compared with the vehicle control group. Notably, the plasma OXT level was markedly elevated after exposed to the chronic restrain stress compared with the vehicle control group (n=8, *P*<0.05;Fig.5D). Down-regulation of OXTR or β-arrestin 2 expression by siRNA transfection markedly inhibited the promotion of migration and invasion of B16 melanoma cells induced by OXT *in vitro* (Fig. 6A-B). Importantly, knockdown of either OXTR or β-arrestin2 in tumor cells significantly inhibited the chronic stress-promoted lung metastasis and extended survival in mice compared with stress group (*n* = 8, *P* < 0.05;Fig. 6C). Immunohistochemistry staining of lung metastasis samples showed the chronic restraint stress up-regulated the ERK1/2 phosphorylation and the expression of OXTR, MMP2 and VEGF in B16 melanoma cells. Strikingly, down-regulation of either OXTR or β-arrestin 2 expression by siRNA transfection significantly diminished phosphorylated ERK1/2 and the production of MMP-2 and VEGF (Fig. 6D).

**Fig. 5:**
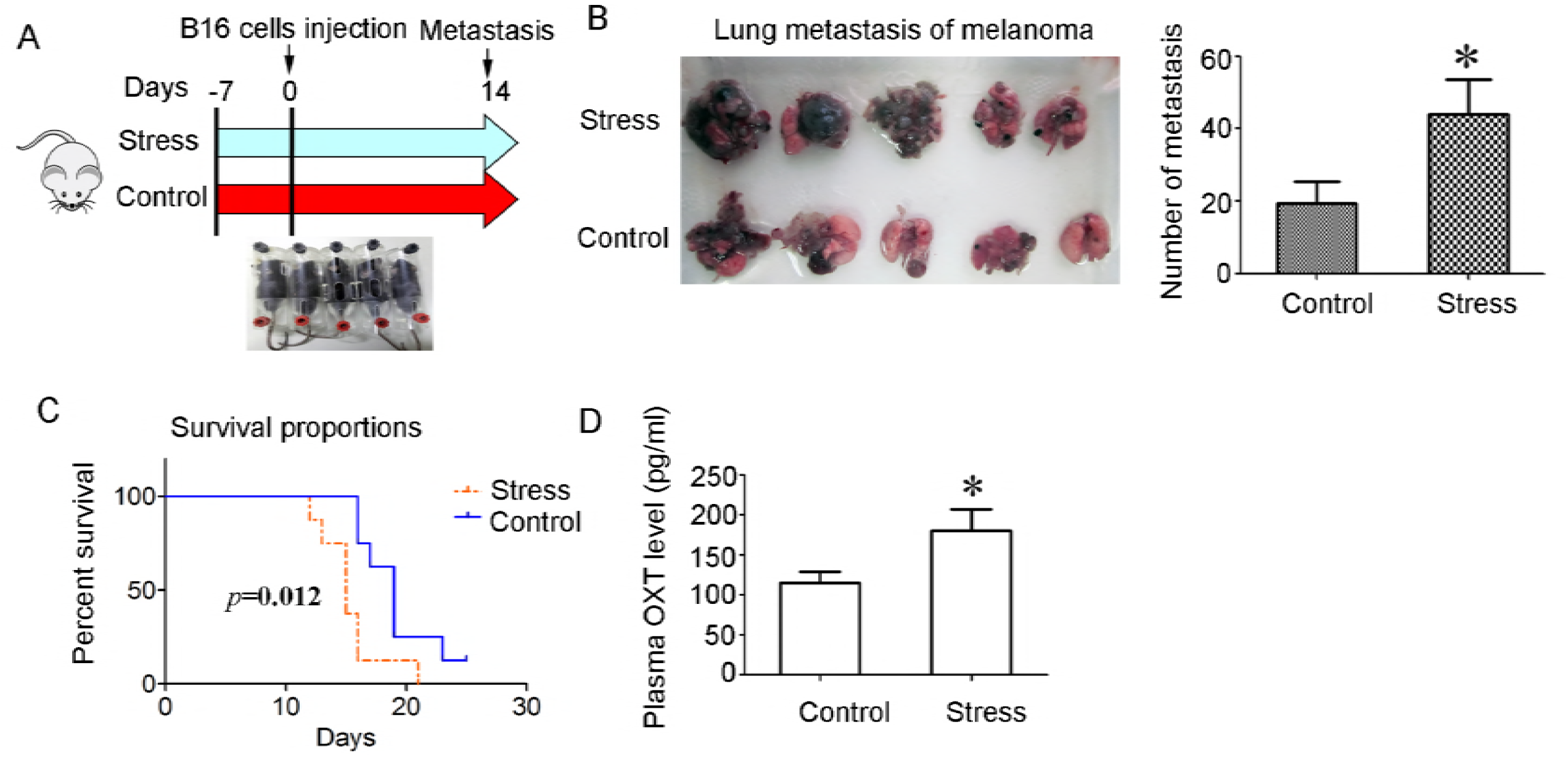
Chronic stress remodels OXT level to promote lung metastasis of melanoma. (A)Schematic representation of the chronic stress paradigm. (B) Representative photograph of pulmonary metastatic foci produced 14 days after intravenous injection of B16 cells. The quantification of the metastatic foci indicated that stress mice developed more metastatic foci on the lung surface than control mice (*n* = 8). \**P* < 0.05. (C) Survival of stress mice and control mice. (D) ELISA of plasma OXT level in control mice and mice subjected to daily restraint stress (*n* = 8). \**P* < 0.05.

**Fig. 6:**
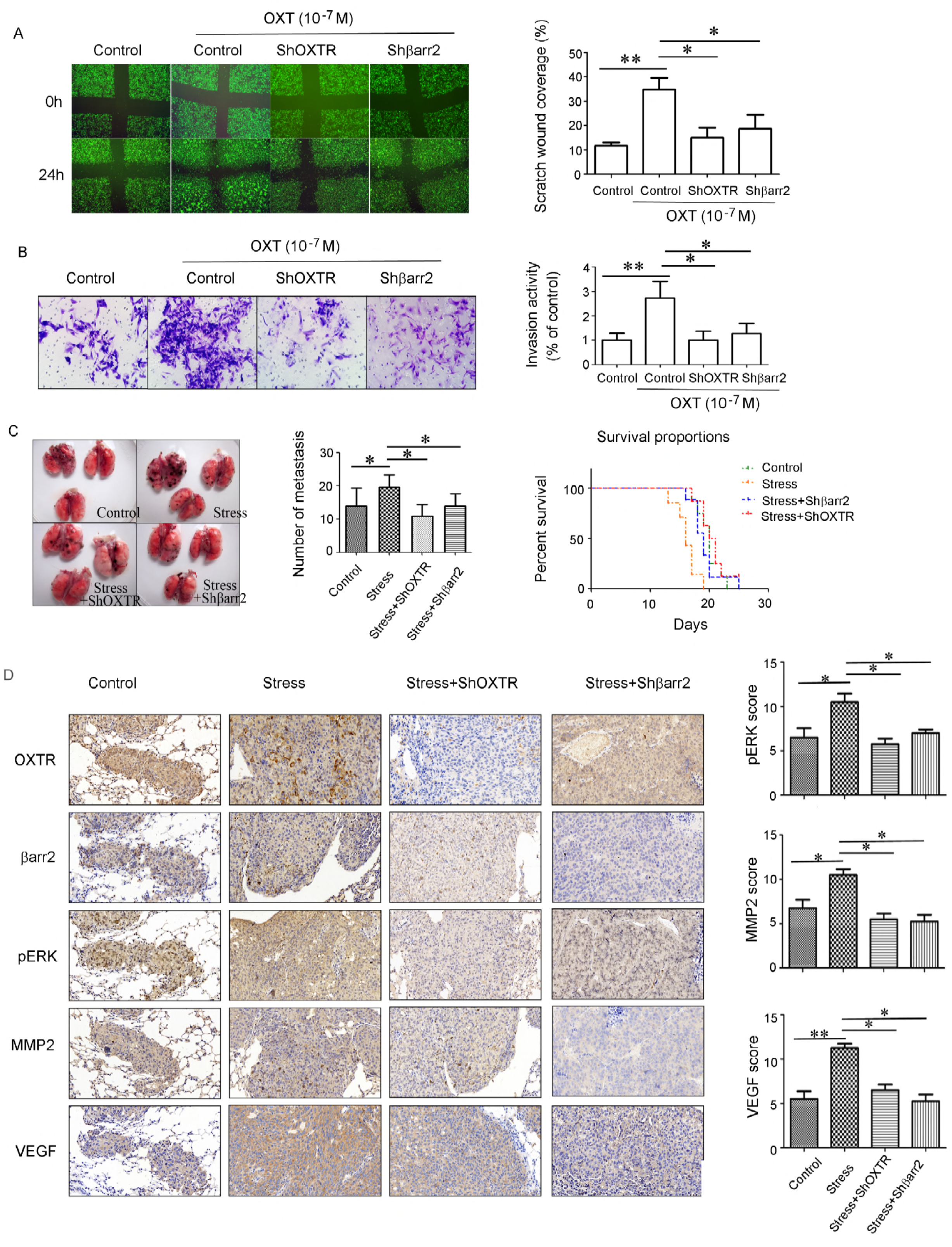
Effect of silencing OXTR or β-arrestin2 on stress-promoted melanoma metastasis *in vivo*. B16 cells were stably transfected with shβarr2, ShOXTR or control. The effect of down-regulation of OXTR or β-arrestin 2 on B16 cells treated with OXT was detected *by in vitro* Wound-healing assay (A) and invasion assay (B). Male mice were subjected to daily restraint stress for 1 week and then injected with B16 cells stably transfected with shβarr2, ShOXTR or control. Two weeks after B16 cell injection, mice were euthanized and the number of nodules were noted. OXTR or β-arrestin2 down-regulation significantly improved the prognosis of chronically stressed tumor bearing animals (*n* =8). The Kaplan-Meier survival curve was depicted (C). Immunohistochemical staining of lung metastasis samples showing the effects of chronic restraint stress on ERK phosphorylation, MMP2 and VEGF (D). OXTR or β-arrestin 2 down-regulation significantly diminished phosphorylated ERK, MMP-2 and VEGF production. All photographs were taken at 400×magnification. The bars in the graphs correspond sequentially to the labeled columns of the images at left. Continuous variables were recorded as mean ± SD. (*n*=3) \**P* < 0.05;\*\**P* < 0.01.

## Discussion

Our results demonstrate that chronic stress promotes melanoma lung metastasis which is mediated by OXT via β-arrestin 2 G protein-independent pathway. It is known that plasma OXT level is increased under stress conditions[26-28]. Our present findings expand the understanding of these processes that regulate melanoma cancer metastasis, indicating a novel strategy for inhibiting metastatic spread through targeted inhibition of OXTR signaling pathway.

### OXTR and its transduction mechanisms

OXT exerts its biological effects through binding to OXTR, which belongs to a family of seven membrane-spanning receptors that transduce signals through G proteins. The coupling of OXTR with Gq results in the activation of PLC, the effector of phosphoinositide signaling system. Several literatures support the conclusion that PLC is the major effector enzyme for transduction of OXT signaling in cellular functions [29, 30]. The oxytocinergic system (OXT and OXTR) exists in various neoplastic cells, where it may stimulate or inhibit cell proliferation. OXTRs are expressed or over-expressed in cells of several tumor types such as breast and endometrial carcinomas [31-34], prostate carcinomas [35], neuroblastomas and glioblastomas [36], osteosarcomas [37], choriocarcinomas [38], and small-cell lung carcinoma [39]. OXTRs were found in the most of ovarian carcinoma tissues [40].

Moreover, all OXTRs in cancer cells are structurally identical with OXTRs in healthy cells and have the same ability to bind OXT [41]. Here we first show that OXTR is expressed in melanoma cells. Importantly, malignant melanoma over-expressed OXTRs. Therefore, *in vitro and in vivo*, OXTRs activation significantly promoted the metastasis of melanoma, but did not promote cell proliferation. Previous studies have shown that OXT inhibits the proliferation of epithelial, nerve and bone derived tumor cells *in vitro*. In turn, OXT was found in tumor cells derived from trophoblasts and endothelial cells to promote cell proliferation [15, 34, 38]. Localization of OXTR within the cell membrane was found to play a critical role in determining the effect of OXT of cell growth. When the OXTR is located outside lipid rafts, OXTR stimulation inhibits cell growth. In contrast, activation of OXTR in lipid rafts stimulates cell growth [42]. Surprisingly, OXT did not exert any effect on proliferation of melanoma cells, the underlying mechanisms of which need to be further investigated. Apart from traditional G-protein dependent signal pathway, β-arrestin mediated G-protein independent pathway has been well studied [43]. In addition to its role in mediating desensitization of GPCRs, β-arrestin has also recently been identified to mediate GPCR-stimulated intracellular signaling [44-49]. Examples of GPCRs that show β-arrestin-mediated signaling are the parathyroid receptor [50], angiotensin II type 1A receptor [49], β1-and β2 adrenergic receptor [47, 48] and vasopressin V2 receptor [51]. Notably, β-arrestin also mediates OXTR signaling [52]. Following agonist stimulation of the OXTR, GPCR kinases phosphorylation of the receptor allows for the recruitment and binding of β-arrestin. β-arrestin recruitment to the OXTR leads to receptor internalization, uncoupling the receptor from G proteins, blocking further signaling. The molecular mechanism by which β-arrestin transduces a signal appears to be through its scaffold function, which has been best characterized for the MAPK pathway [53].

### OXTR/β-arrestin 2-ERK signal pathway in melanoma metastasis

To determine whether the β-arrestin-dependent manner mediates OXT-induced melanoma cells migration and metastasis, we performed β-arrestin loss-of-function experiments *in vitro* and *in vivo*. B16 melanoma cells transfected with siRNA directed against β-arrestin 2 demonstrate decreased ERK phosphorylation following OXT treatment. Because β-arrestin1 is not located in melanoma cells, we draw a conclusion that only β-arrestin-2 is necessary for β-arrestin-dependent ERK signaling from the OXTR activation in melanoma cells. The present studies first revealed β-arrestin2 is involved in OXT-stimulated melanoma metastasis. However, our results does not rule out the possibility that traditional G-protein dependent pathway also involves in OXT-induced ERK activation. An important implication of this work is that the conformations of OXTR induced by OXT is competent to activate ERK1/2 via β-arrestin 2, but not able to activate G proteins.

Metastasis is a major cause of cancer morbidity and mortality. Multiple host cell types contribute to metastasis and recent studies have focused on host microenvironment as a new target for anti-metastasis therapy. Neuroendocrine kinetics has the potential to regulate the expression of genes of various cell types in tumor microenvironment. Interestingly, our current study suggests that in mouse melanoma models, inhibiting stress promotes melanoma metastasis and inhibits survival, consistent with previous studies [9, 10]. Strikingly, genomic knockdown of OXTR or β-arrestin 2 inhibited or even cured the melanoma metastasis, respectively.

## Conclusions

These findings suggest that increased OXT under stress may contribute to stress-related melanoma metastases. Therefore, as far as we know, it first provides a possible explanation for promoting melanoma metastasis under these conditions. Clinically, current data suggest that localized targeting of tumor-associated OXTR signal pathway (for example, with antagonists or anti-oxtr antibodies) may constitute a new auxiliary strategy to minimize melanoma metastasis.

## Declarations

### Ethics approval and consent to participate

A total of 15 pairs of malignant melanoma tissues and adjacent normal tissues were collected from Shandong Cancer Hospital Affiliated to Shandong University after written informed consents were obtained. The study protocol was approved by the Ethics Committees of Shandong Cancer Hospital Affiliated to Shandong University, Jinan 250117, China. Male C57/B6 mice (6-8 week of age) were obtained from Shandong University Animal Centre. The experimental procedures were approved by the Animal Ethic Committee at Shandong University.

### Consent for publication

Not applicable

### Availability of data and material

Data sharing not applicable to this article as no datasets were generated or analysed during the current study.

### Competing interests

The authors declare no potential conflicts of interest

### Funding

This work was supported by grants from the National Natural Science Foundation of China (31571183), Open Project of National Research Center for Assisted Reproductive Technology and Reproductive Genetics, Shandong University and Shandong Key Research and Development Program (2017GSF218032).

### Authors’ contributions

HJ and NL performed the experiments and took active part in the writing of the manuscript. DC, QS and DL performed Western Blot and immunochemistry for OXTR expression. YL performed ELISA for plasma OXT. JL, BB, YL and YY collected clinical MM samples. NL performed the statistical analyses. XC revised the manuscript. JL was responsible for the design of the study, interpretation of data and writing of the manuscript.

## Acknowledgements

Not applicable

---

**Fig. S1 Silencing either OXTR or β-arrestin 2 largely suppressed the expression of OXTR or β-arrestin 2 in B16 melanoma cells.**

The representative photograph of control siRNA (CTL siRNA), OXTR siRNA (A) or β-arrestin 2 siRNA (B).

**Fig. S2 The expression of β-arrestin 1 in B16 melanoma cells** B16 cells stably transfected with shβarr2, ShOXTR or control were injected in mice 1 week after chronic restraint stress. Two weeks later, the animals were killed by cervical dislocation. Immunohistochemical staining of lung metastasis samples for β-arrestin 1. The B16 melanoma cells did not express β-arrestin 1.

**Supplemental table 1. The clinicopathological information of malignant melanoma**

